# *Galleria mellonella* as a novel eco-friendly in vivo approach for the assessment of the toxicity of medicinal plants

**DOI:** 10.1101/2021.07.06.451318

**Authors:** Mbarga Manga Joseph Arsene, Podoprigora Irina Viktorovna, Anyutoulou Kitio Linda Davares

## Abstract

The evaluation of medicinal plants’ toxicity is a prerequisite prior their usage. The vertebrate models used for this purpose are often the object of ethical consideration. Though invertebrate models including *Galleria mellonella* have shown their ability to be used to assess various products’ toxicity, to our knowledge, *G. mellonella* has never been exploited to determine the toxicity of medicinal plants. In this study, the toxicity of hydroalcoholic and aqueous extracts of seven (7) Cameroonian medicinal plants namely leaves of *Cymbopogon citratus* (DC.) Stapf, *Moringa oleifera* Lam and *Vernonia amygdalina* Delile; barks of *Cinchona officinalis* and *Enantia chloranta* Oliv; barks and seeds of *Garcinia lucida* Vesque and leaves and seeds of *Azadirachta indica* (Neem) were evaluated using the larval form of the Greater Wax Moth (*Galleria mellonella*). The median lethal doses (LD50), 90% lethal doses (LD90) and 100% lethal doses were successfully determined using the spline cubic survival curves and equations from the data obtained on the survival rate of *G. mellonella* 24h after the injection with the extracts. The LD50 values varied from 3.90 g/kg bw to >166.67 g/kg bw and the pattern of toxicity observed was in accordance with previous investigations on the plant materials concerned. The results obtained in this study suggest that *G. mellonella* can be used as a sensitive, reliable, and robust eco-friendly model to gauge the toxicity of medicinal plants. Thus, avoid the sacrifice of vertebrate models often used for this purpose to limit ethical concerns.

## 1. Introduction

Medicinal plants have been used for millennia to treat various pathologies including respiratory tract infections, urinary tract infections, sepsis, diabetes, erectile insufficiency, diarrhea, fevers and as a stabilizer of physiological functions among many others. In this study, the herbal medicines used were leaves of *Cymbopogon citratus* (DC.) Stapf, *Moringa oleifera* Lam and *Vernonia amygdalina* Delile; barks of *Cinchona officinalis* and *Enantia chloranta* Oliv; barks and seeds of *Garcinia lucida* Vesque and Leaves and seeds of *Azadirachta indica* (Neem). These medicinal plants are among the most famous in Cameroon.

*Cinchona officinalis* is a shrub of the Rubiaceae family whose bark is recognized for its very bitter taste and its composition in alkaloids such as quinine, dihydroquinine, cinchonidine, epiquinine, quinidine, dihydroquinidine, cinchonine and epiquinidine [1,2]. In addition to its antimalarial properties, *C. officinalis* has also been used on a smaller scale as a treatment for goiter, Meniere’s disease (an inner ear disorder), varicose veins, and in obstetrics for its action on uterine muscles [2].

*Garcinia lucida* Vesque (*G. Lucida)* is also a well-known herbal medicine whose properties and composition have been studied. The seed, fruit and bark are the most commonly used parts in traditional medicine and food [3]. Used in fermentation of palm tree or raffia wine [4], the bark and seeds have been found useful in the treatment of gastric and gynecological infections, diarrheas, cure for snake bites as well as an antidote against poison [3]. In addition, a recent study conducted by Sonfack et al. [5] revealed that the aqueous extract from the stem bark of *G. lucida* exhibits cardioprotective and nephroprotective effects in adenine-induced chronic kidney disease in rats. This plant is also believed to possess some aphrodisiac properties and also used to hunt ghosts [6]. Investigations on the chemical composition of *G. lucida* have shown that it contains the terpenoids and xanthones products such as 1,2-dihydroxy-xanthone, 1-hydroxy-2-methoxyxanthone, putranjivic acid, methyl putranjivate and friedelin [3,7].

*Enantia chloranta* Oliv (*E. chloranta*) is a plant belonging to the Annonaceae family with multiple uses in traditional medicines. Also called Epoue (Baka), Peye (Badjoue), and Nfol (Bulu), it is widely spread along Sub-Saharan Africa [8]. In Cameroon, *E. chloranta* is used in the management of various infections including malaria, typhoid fever, jaundice, dysentery, wounds, high blood pressure, urinary infection, leprosy spots and convulsions. Its yellow, dried, or fresh bark has been reported to have antiviral [9], antifungal [10] and antibacterial properties [8,10]. Other studies have reported its use as an antioxidant and antipyretic [11]. Investigations into its composition have revealed that *E. chloranta* contains a wide variety of biologically active compounds, the most important of which are alkaloids such as protoberberines (berberine, canadine, palmatine, jatrorrhizine, columbamine and pseudocolumbamine), phenanthrene alkaloids (atherosperminine and argentinine) and aporphines (7-hydroxydehydronuciferine and 7-hydroxydehydronornuciferine) [11].

*Azadirachta indica* (*A. indica)* known as Neem is a monoecious tree of the Meliaceae family whose oil produced from its seeds is widely used for its medicinal properties in the northern part of Cameroon. It is known that compounds in Neem extracts have anti-inflammatory, anti-hyperglycaemic, anti-carcinogenic, antimicrobial, immune-modulator, anti-mutagenic, antioxidant, anti-ulcer and anti-viral effects [12,13] Recent studies by Baildya et al. [13] found 19 compounds from this plant which may be used as anti-COVID-19. In addition, Azadirachtin, one of its major constituents, gives it insecticidal properties which acts by blocking metamorphosis from the larval to adult stage, and paralyzes the digestive tract of the larva [14,15]. The different parts of *A. indica* have been widely used in recent years in the green synthesis of nanoparticles such as silver nanoparticles (AgNPs), Zinc oxide nanoparticles (ZnONPs), gold (AuNPs) and copper nanoparticles (CuNPs) [16–19].

*Moringa oleifera* Lam (*M. oleifera*) is both an edible and a medicinal plant. Every part of the plant, from the leaves to the roots, has been reported to possess potential health benefits [20]. This plant belonging to the Moringaceae family contains a profile of important minerals, and is a good source of protein, vitamins, β-carotene, amino acids and various phenolic compounds [21]. *M. oleifera* provides a rich and rare combination of zeatin, quercetin, β-sitosterol, caffeoylquinic acid and kaempferol [20,21]. Besides its nutritional properties, *M. oleifera* is traditionally used to treat skin infection, asthma, diabetes, diarrhea, arthritis, inflammation, cough, fever, and headache. It has also been reported to have Antioxidant, anti-inflammatory, antitumor, antimicrobial, hepatoprotective and anti-arthritic properties [22–24].

Like *M. oleifera, Vernonia amygdalina* Delile (*V. amygdalina*) is both an edible and a medicinal plant. *V. amygdalina* belongs to the family Asteraceae and is called bitter leaf in English because of its bitter taste. In Cameroon, *V. amygdalina* is known under the name of Ndolè and is included in the composition of the national popular dish which bears the same name. The presence of polyphenols, vitamins and mineral salts make the plant useful in human diets [25,26]. *V. amygdalina* is very rich in phytochemicals such as saponins, sesquiterpenes, flavonoids, and steroid glycosides (vernosides), lactones [26] and several studies reported that it has anticancer and antitumor activity [27,28]; antihepatotoxic activity [29]; hypoglycemic activity [26]; antibacterial activity [30]; anti-inflammatory [31] as well as antioxidant property [32].

*Cymbopogon citratus* (DC.) Stapf (*C. citratus*) or lemongrass is an herbaceous plant of the Poaceae family growing in humid tropical areas which is mainly cultivated for its stems and leaves whose infusion gives a tea with strong lemony odor due to its high content of the aldehyde citral [33]. Muala et al. [34]. reported that lemongrass is rich in minerals, vitamins, macronutrients (including carbohydrate, protein, and small amounts of fat) and its leaves are a good source of various bioactive compounds such as alkaloids, terpenoids, flavonoids, phenols, saponins and tannins that confer *C. citratus* leaves pharmacological properties such as anti-cancer, antihypertensive, anti-mutagenicity, anti-diabetic, antioxidant, anxiolytic, anti-nociceptive and anti-fungi. *C. citratus* is also commonly used in folk medicine for treatment of nervous and gastrointestinal disturbances, and as antispasmodic, analgesic, anti-inflammatory, anti-pyretic, diuretic, and sedative properties [35].

In general, medicinal plants constitute a source of new biologically active molecules that are economically accessible to deal with the emergence of phenomena of resistance of germs to chemical molecules [36]. The common point of all these medicinal plants (and all the molecules that result from them) that can potentially be used in the treatment of various pathologies is the evaluation of their toxicity. Indeed, toxicity testing in rodents is an important prerequisite to the use of compounds in man [37]. However, trials in rats and mice are expensive, not eco-friendly and there are ethical considerations [37]. In this context, invertebrate models appear as a very promising alternative and have seen a gain in popularity in recent years [37–41]. The most used invertebrate models for toxicity trials among others are *Artemia salina, Daphnia magna*, *Drosophila melanogaster*, soil invertebrate *Folsomia candida*, the nematode *Caenorhabditis elegans*, and the Greater Wax Moth *Galleria mellonella* [38,42–44]. Among all these in vivo insects’ models, *G. mellonella* has several attractive benefits due to its practicality, in particular its low cost, ease of acquisition and handling (no specialist training or equipment is required), ability to survive at 37°C (unlike many other non-mammalian hosts), they are large enough (2 cm in length and weigh approximately 250 mg) for accurate dosing (unlike commonly used invertebrate models such as *C. elegans* and *D. melanogaster*) which make it possible to handle individual larvae and administer test compounds directly into each individual (which is not always possible with other smaller invertebrate models) and enhances the reproducibility of assays conducted using this model [37,38,45].

The aim of this study was to demonstrate the potential of *G. mellonella* larvae as a simple, inexpensive, and rapid model for the evaluation of the toxicity of herbal medicines such as *Cinchona officinalis, Garcinia lucida* Vesque, *Enantia chloranta* Oliv, *Azadirachta indica* (Neem), *Moringa oleifera* Lam, *Vernonia amygdalina* Delile and *Cymbopogon citratus*. To our knowledge, this study presents the first use of *G. mellonella* for an in vivo LD50 study of medicinal plants.

## 2. Material and method

### 2.1. Vegetal material

The vegetal materials used in this study were leaves of *Cymbopogon citratus* (DC.) Stapf, *Moringa oleifera* Lam and *Vernonia amygdalina* Delile; barks of *Cinchona officinalis* and *Enantia chloranta* Oliv; barks and seeds of *Garcinia lucida* Vesque and Leaves and seeds of *Azadirachta indica* (Neem). These plants were chosen because they are renowned for their use in traditional medicine in Cameroon and because of data existing in the literature on their effectiveness and their composition. All the plants were collected in September 2020 in different regions of Cameroon. *V. amygdalina and C. citratus* was collected in Nlobison II, Centre region of Cameroon (VJ5J+C3 Yaounde, Cameroon); *C. officinalis*, *E. chloranta and G. lucida* were bought in the Nkoabang Market (VH7M+FJ Yaoundé, Cameroun); *M. oleifera* was bought in the Dang Market of Adamaoua region (CHH5+67 Ngaoundéré, Cameroun) and *A. indica* was bought in the North region (7CX2+X4 Garoua, Cameroun). The collected plants were dried at room temperature in the shade for 7 days then packaged in hermetically sealed plastics and additional packaging was done to facilitate shipment to Russia in December 2021 where they were received by the Laboratory of Microbiology, Faculty of Medicine of the RUDN University. Plants were grinded and the powders with particle sizes lower than 1 mm were stored in a sterile airtight container until further use.

### 2.2. Galleria *mellonella*

*G. mellonella* larvae were obtained commercially from https://ecobaits.ru/ (ECO BAITS, Moscow, Russia) and stored at 15°C prior to use. Dead larvae and those with dark spots or showing signs of melanisation were discarded.

### 2.3. Extraction of active compounds

Hydroalcoholic solution (80%, v/v) and distilled water was used because they have been reported to be an efficient solvent for the extraction of bioactive compounds in the medicinal plants used in this study. As we described in our previous investigation [46], fifty grams (50g) of vegetal material was weighed and added to 450 ml of the solvent in separate conical flasks. The flasks were covered tightly and were shaken at 200 rpm for 24h and 25°C in a shaker incubator (Heidolph Inkubator 1000 coupled with Heidolph Unimax 1010, Germany). The mixtures were then filtered by vacuum filtration, using Whatman filter paper № 1 then concentrated at 40°C in rotary evaporator (IKA RV8) equipped with a water bath IKA HB10 (IKA Werke, Staufen, Germany) and a vacuum pumping unit IKA MVP10 (IKA Werke, Staufen, Germany). To avoid losses, the extracts were collected when the volumes were small enough and placed in petri dishes previously weighed and then incubated open at 40°C until complete evaporation. The final dried crude extracts were weighed. Extraction volume and mass yield were determined using the following formulas:

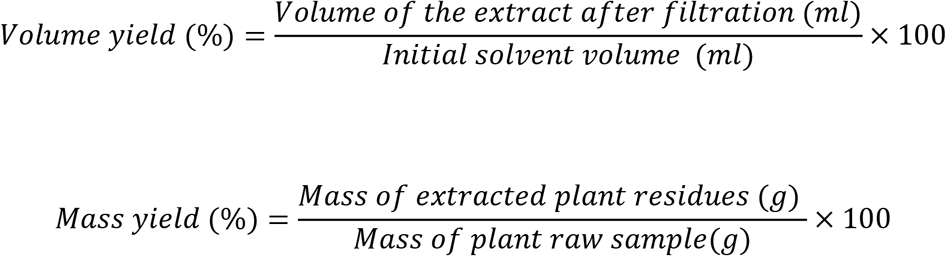

### 2.4. Stock solutions

1g of each crude extract was dissolved in 5 mL of sterile phosphate-buffered saline (PBS) to obtain the concentration of 200mg/ml. These solutions were sterilized by microfiltration (0.22μm) and dilutions were performed in sterile PBS to obtain concentrations of 150, 100, 50, 25, 12.5 and 5mg/ml, the sterile PBS plant extract free were used as control. All the 8 tests solution were stored at 4°C.

### 2.5. Toxicity screen

The larvae were weighed and only those from 0.2 to 0.5g were retained. For each concentration of extract, 3 groups of twenty (20) randomly selected *G. mellonella* larvae were used (Figure 1-A). 20μl of each dilution were injected using a 0.3 ml Terumo® Myjector® U-100 insulin syringe (VWR, Russia) through the base of the last left proleg as described by Megaw et al [38] and shown in Figure 1-B. Control groups of 10 larvae injected with 20 μl sterile PBS were also included. After 24h of incubation in the dark at 37°C, larvae were examined for mortality, and were considered dead if they were unmoving, failed to reorient themselves when placed on their backs, and failed to respond to stimuli as indicated by Megaw et al [38]. Percentage survival was plotted as a function of concentration for each of the plants extract using Spline cubic model in the statistical software XLSTAT 2020 (Addinsof Inc., New York, USA) and median lethal dose (LD50), 90% lethal dose (LD90) and the 100% lethal dose (LD100) values expressed in mg/ml were calculated using each specific spline cubic equation or spline cubic curves obtained. Spline curves were used when the fit accuracy of the spline equation was less than 90%). The mean weight of each group of 20 larvae was used to extrapolate the LD50, LD90 and LD100 values for each plant extract in g/kg using the formulas:

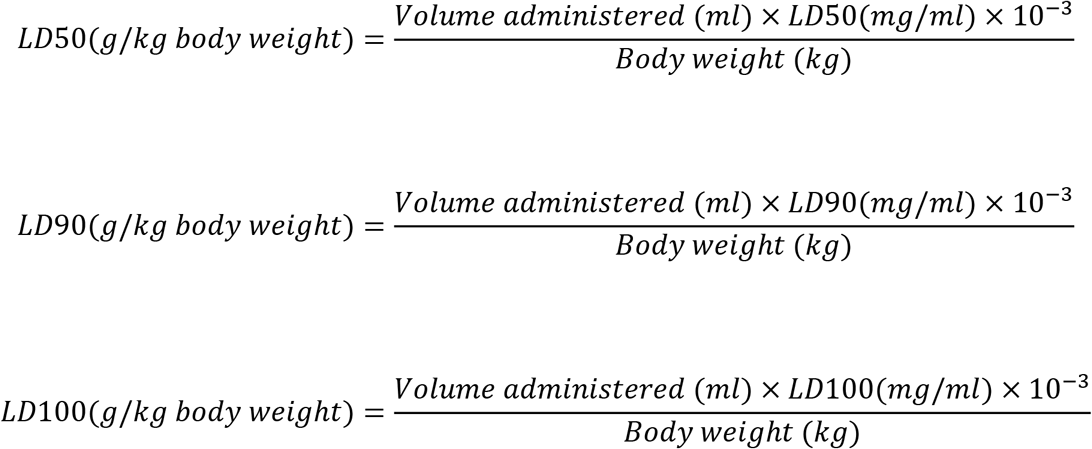

**Figure 1:**
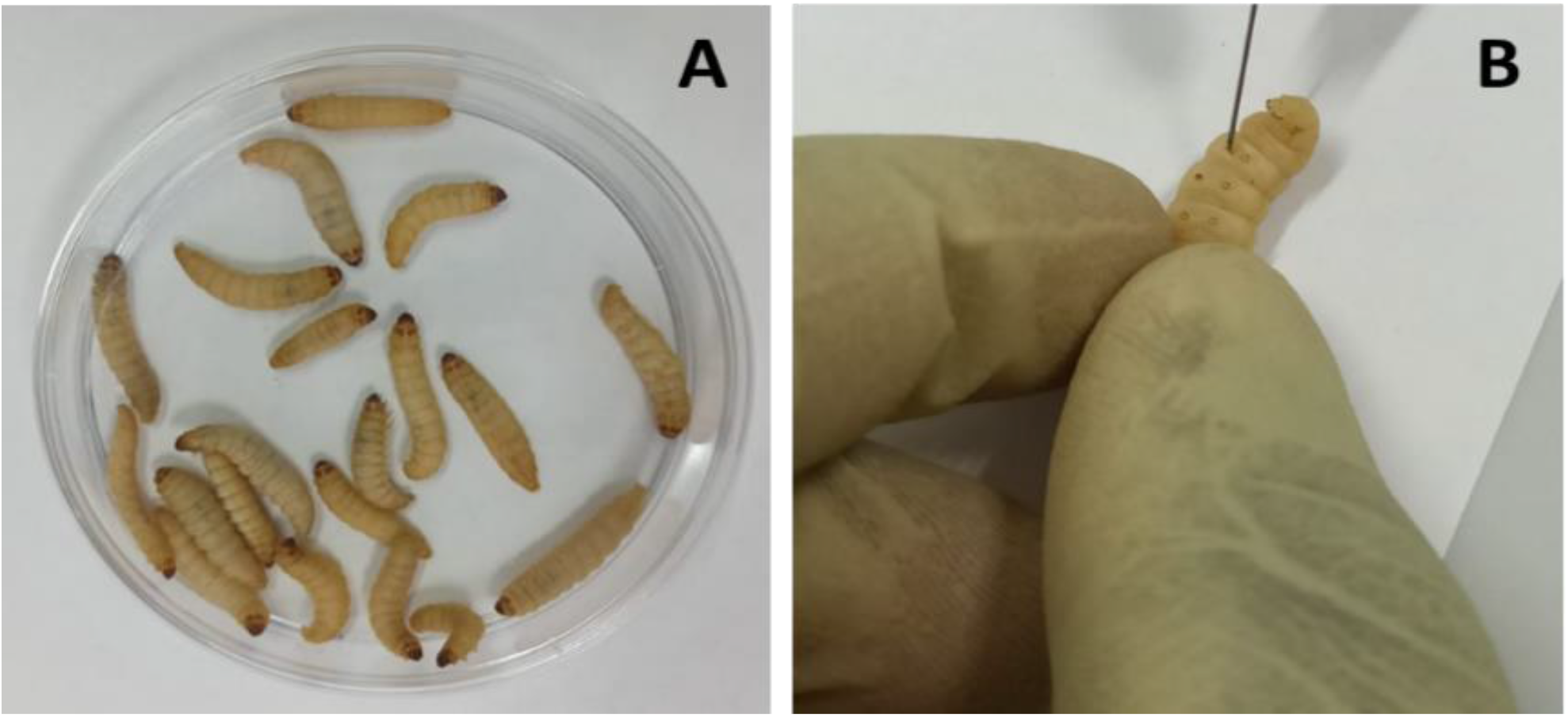
(A)Group of 20 larvae and (B) Injection of *G. mellonella* larvae with plant extract through the last left proleg.

## 3. Results and Discussion

The extract yields obtained for the 9 samples from our 7 medicinal plants using hydroalcoholic solution and distilled water are recorded in Table 1. As observed in Table 1, the highest volume yields were observed with hydroalcoholic extracts while the highest mass yield were obtained with aqueous leaves extract of *M. oleifera* (21.2%), *C. citratus* (17.3%), *A. indica* (15.1%) and *V. amygdalina* (14.0%). The highest mass yield with hydroalcoholic solution was obtained with *E. Chloranta* bark (12.4%). Extraction with hydroalcoholic solution was less effective in *A. indica* seed and *M. oleifera* leaves with mass yield of 5.8% and 7.3% respectively. Most of the studies that used different solvents, whether for the plants used in our study or for other plants, it was reported that water had a better extraction rate than other solvents such as ethanol or methanol [46,47]. Mouafo et al [47] reported that this difference is mainly attributed to the polarity of the solvents and therefore to the large proportions of water-soluble compounds present in the plant. However, high yields of phytoconstituents obtained does not necessarily imply a high biological activity [47]. We observed this in our study through the lethal doses (LD50, LD90 and LD100) determined for each of the extracts of the medicinal plants tested. Indeed, as shown in Table 3, LD50, LD90 and LD100 values vary considerably from one plant to another but above all, we noticed that for all medicinal plants, the lethal doses of hydroalcoholic extracts were lower than those of aqueous extracts. This implies that hydroalcoholic extracts contain more active compounds and are therefore more toxic than aqueous extracts because, as reported by Situmorang et al [48], low LD50 values are associated with high toxicity (high amount of active compounds) and conversely, high LD50 values are associated with low toxicity (low amount of active compounds).

**Table 1:**
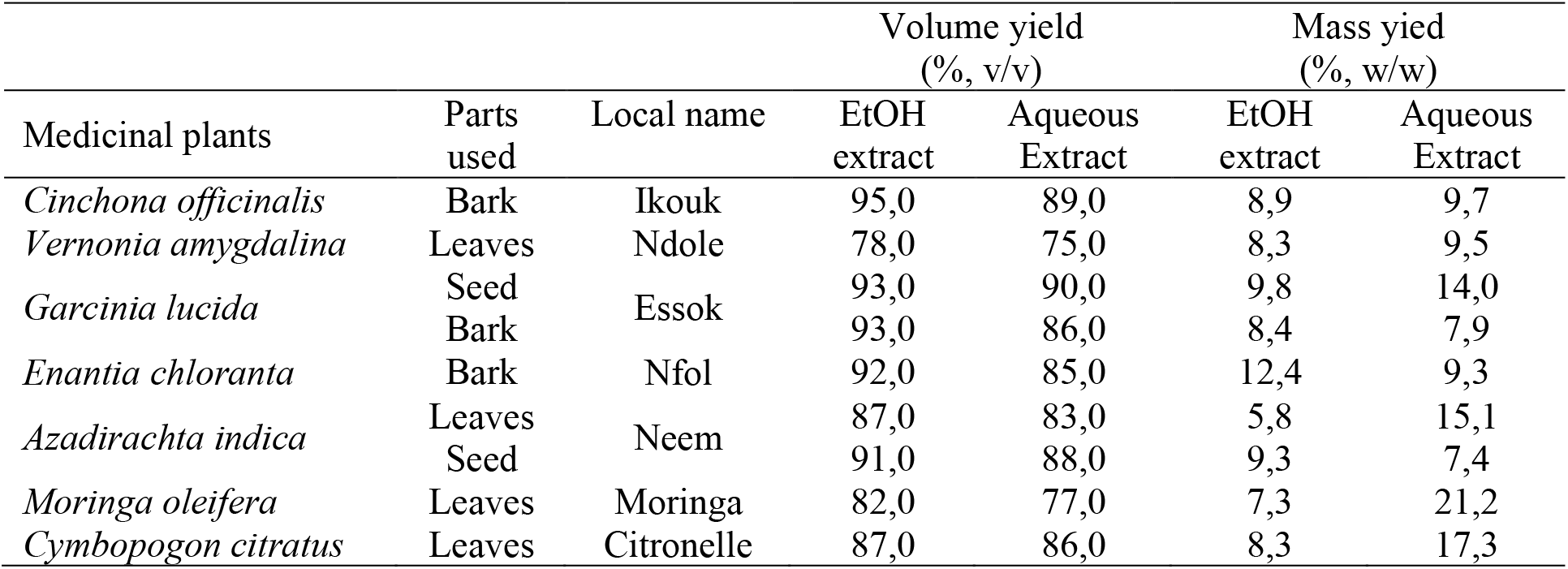
Extract yields (%) obtained from the seven plants with distilled water and hydroalcoholic solutions (EtOH 80%)

It is important to remember that the LD values were obtained using the Greater Wax Moth (*Galleria mellonella*) as a toxicity assessment model. After 24 hours incubation, survival data for each concentration of each plant material was used to plot spline cubic survival curves (examples of which are shown in Figure 2) and to obtain the spline cubic equations (examples of which are presented in Table 2). The spline cubic equations and curves of all the other extracts can be found in supplement material (Supplement 1: Table 1 and Figure 1). As expected, no deaths were observed in larvae injected with 20μl of Phosphate-buffered saline (PBS). However, as shown in Table 3, the LD50 (mg/ml) of the extracts tested varied from 4,87 (3,90 g / kg bw (body weight)) to >200 (> 166,67 g/kg bw), the LD90 (mg / ml) from 25,00 (18.52 g/kg bw) to >200 (> 181,82 g/kg bw) and LD100 (mg/ml) from 45,00 (40.91 g/kg bw) to > 200 (>181.82 g/kg bw). This difference between the LD values confirms the good sensitivity of *G. mellonella*, which was reported as changing depending on the substances tested [38]. The extracts with the highest toxicity were *Cinchona officinalis* hydroalcoholic bark extract (LD50 = 3,90 g/kg bw) followed by *Azadirachta indica* hydroalcoholic seed extract (LD50 = 4,81 g/kg bw) and *C. officinalis* aquous bark extract (LD50 = 8,66 g/kg bw). However, other plant materials such as water extract of leaves of *C. citratus*, *M. oleifera* and *V. amygdalina* exerted the lowest toxicity on the greater wax moth with LD50 (mg/ml) respectively of 149,08; 165,40 and > 200.

**Table 2:** Cubic spline model used to determine the LD50, LD90 and LD100 of *E. chloranta bark* hydroalcoholic extract (HAE), *C. citratus* leaves HAE, *A. indica seed* HAE and *G. lucida* bark HAE.

| Plant material | Spline intervals | t<sup>3</sup> | t<sup>2</sup> | t | Constant | R<sup>2</sup> |
| --- | --- | --- | --- | --- | --- | --- |
| *E. chloranta* bark HAE | [0;5] | -0,005 | 0,000 | 0,119 | 100,000 | 0,92 |
|  | [5;12.5] | 0,002 | -0,072 | -0,239 | 100,000 |  |
|  | [12.5;25] | 0,001 | -0,028 | -0,986 | 95,000 |  |
|  | [25;50] | 0,000 | 0,005 | -1,278 | 80,000 |  |
|  | [50;100] | 0,000 | 0,000 | -1,163 | 50,000 |  |
|  | [100;200] | 0,000 | 0,010 | -0,668 | 0,000 |  |
| *C. citratus* leaves HAE | [0;5] | 0,003 | 0,000 | -0,067 | 100,000 | 0,88 |
|  | [5;12.5] | -0,008 | 0,040 | 0,133 | 100,000 |  |
|  | [12.5;25] | 0,004 | -0,133 | -0,567 | 100,000 |  |
|  | [25;50] | 0,000 | 0,019 | -2,000 | 80,000 |  |
|  | [50;100] | 0,000 | 0,011 | -1,264 | 40,000 |  |
|  | [100;200] | 0,000 | 0,006 | -0,413 | 0,000 |  |
| *A. indica* seed HAE | [0;5] | 0,038 | 0,000 | -9,951 | 100,000 | 0,69 |
|  | [5;12.5] | -0,021 | 0,570 | -7,099 | 55,000 |  |
|  | [12.5;25] | -0,002 | 0,099 | -2,080 | 25,000 |  |
|  | [25;50] | 0,000 | 0,013 | -0,676 | 10,000 |  |
|  | [50;100] | 0,000 | 0,006 | -0,186 | 0,000 |  |
|  | [100;200] | 0,000 | -0,001 | 0,068 | 0,000 |  |
| *G. lucida* bark HAE | [0;5] | -0,017 | 0,000 | -0,584 | 100,000 | 0,85 |
|  | [5;12.5] | 0,018 | -0,249 | -1,831 | 95,000 |  |
|  | [12.5;25] | -0,005 | 0,165 | -2,468 | 75,000 |  |
|  | [25;50] | 0,001 | -0,025 | -0,721 | 60,000 |  |
|  | [50;100] | 0,000 | 0,016 | -0,937 | 35,000 |  |
|  | [100;200] | 0,000 | 0,000 | -0,137 | 15,000 |  |

**Table 3:**
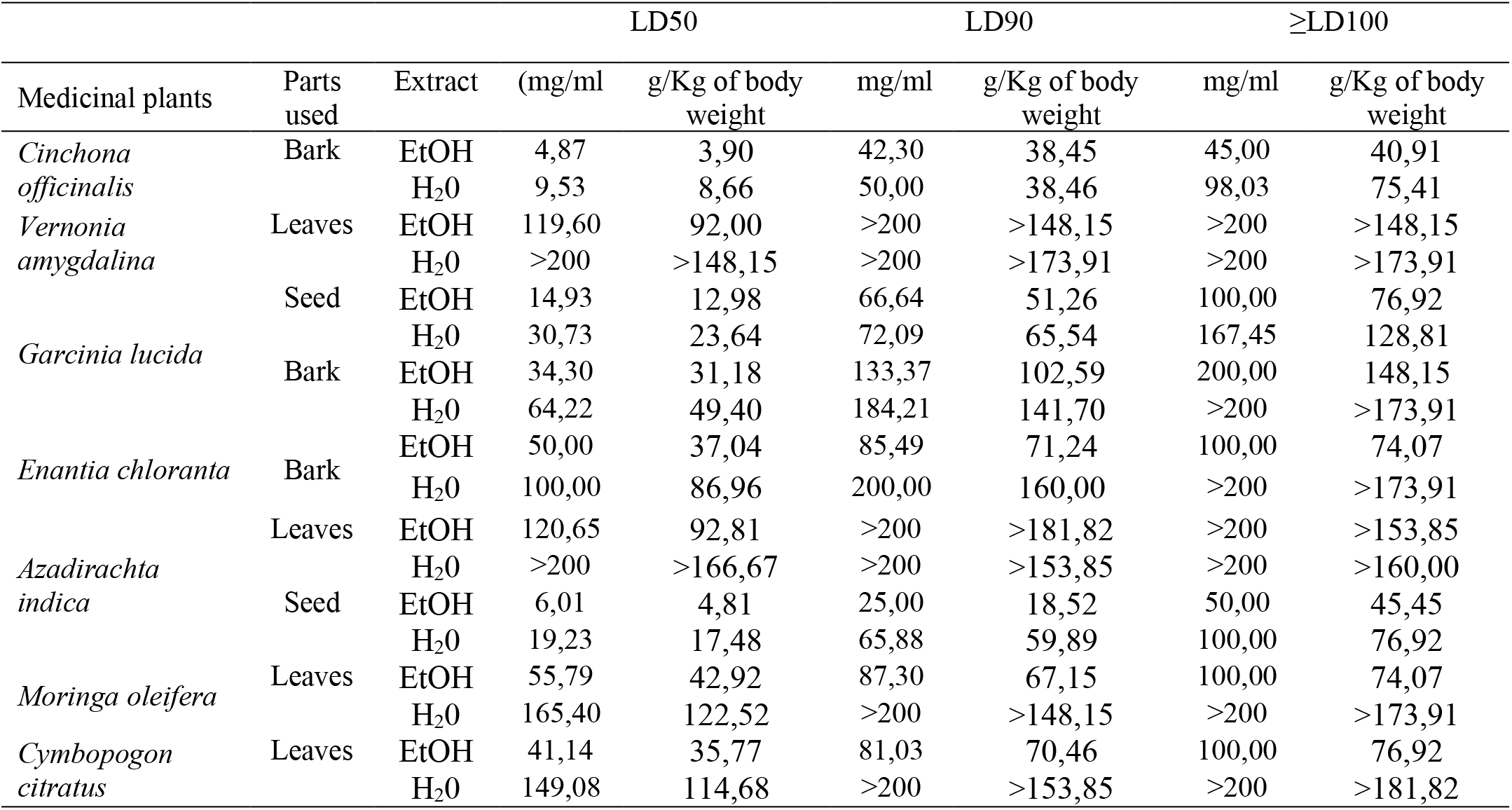
LD50, LD90 and LD100 values for the medicinal plants tested on *G. mellonella* model.

The high toxicity of both hydroalcoholic and aqueous extract of *C. officinalis* bark can be attributed to its composition having abundant quinine and its derivatives (dihydroquinine, cinchonidine, epiquinin, quinidine, dihydroquinidine, cinchonine and epiquinidine) [1,2] which are slightly soluble in water and soluble in ethanol [49]. Indeed, Quinine, C_20_H_24_N_2_O_2_, (8S, 9R) -6′-methoxycinchonana-9ol;(αR)-α-(6-methoxy-4-quinoyl)-α-[(2S, 4S, 5R)-(5-vinylquinuclidin-20yl)] methanol, is the most important alkaloid occurring in Cinchona [50]. We did not find any study in the literature evaluating the toxicity of extracts of *C. officinalis,* but quinine was reported to have an LD50 of 1,392, 0,660 and 0,641 g/kg bw when administered orally to rats, mice, and rabbits respectively [51]. These values are lower than our findings probably because the results reported by ECHA [51] were those of pure quinine while our study focused on extracts with heterogeneous composition. Furthermore, and interestingly, we observed that some larvae exposed to concentrations greater than the LD100 of hydroalcoholic extracts of *C. officinalis* bark exhibited localized melanizations whereas larvae injected by lower concentrations had slight and uniform melanization (Figure 3). These results are similar to those obtained by Megaw et al. [38] who used the *G. mellonella* model to test the toxicity of 1-alkyl-3-methylimidazolium chloride ionic liquids. Megaw et al [38] reported that melanization is a common component of the insect immune response, which occurs as a result of stress or infection, and leads to the larvae changing from cream-colored to dark brown or black. The two types of melanization in our study (Fig 3b and 3c) can be explained by the fact that the high concentrations of the extracts are poorly distributed within the larvae unlike the low concentrations.

**Figure 3:**
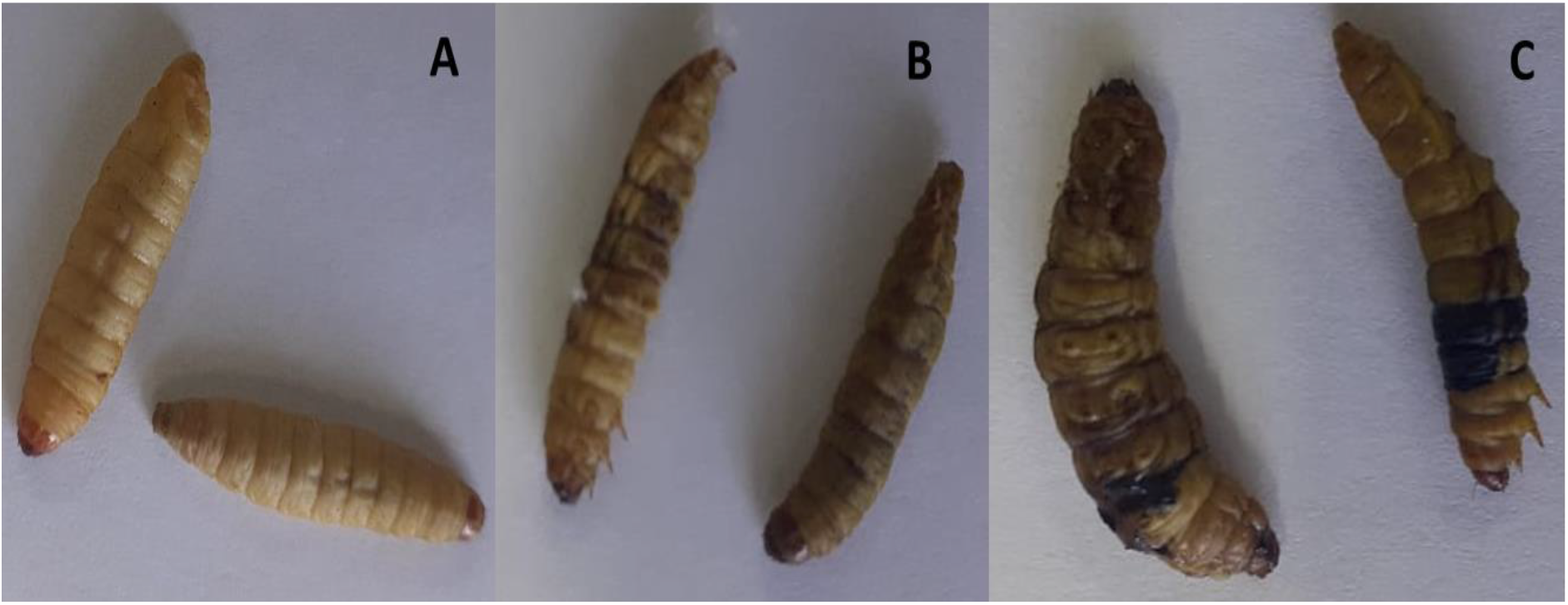
Healthy G. mellonella larvae (A); larvae showing slight and uniform melanisation (B); and larvae showing localised melanisation following injection with a LD100 of Cinchona officinalis hydroalcoholic bark extract.

In addition, similarly to our study, seeds of *A. indica* have been reported to have high toxicity both in animal models [52] and in insects [53] due to phytochemicals that it contains, including azadirachtin, which is used in the composition of certain natural insecticides [14]. This substance acts mainly by blocking metamorphosis from the larval to adult stage and paralyzes the digestive tract of the larva [14,15]. Azadirachtin has been reported to be sparingly soluble in water and freely soluble in polar organic solvents such as methanol, ethanol, and ethyl acetate [54], which could explain the large difference between the LD50s of the. aqueous extract and the hydroalcoholic extract of *A. indica* seed which were respectively 19,01 mg/ml and 6,01 mg/ml. However, both hydroalcoholic and aqueous extracts from the leaves of *A. indica* have demonstrated more than 10 times less toxicity than the seeds, possibly due to their different phytoconstituent composition.

According to the classification of Gosselin, Smith and Hodge [55], biologically active compounds can be super toxic (LD50 <5 mg/kg bw), extremely toxic (LD50 ∈ [5-50 mg/kg bw]), very toxic (LD50 ∈ [50-500 mg/kg bw]), moderately toxic (LD50 ∈ [0,5-5 g/kg bw]), slightly toxic (LD50 ∈ [5-15 g/kg]) and practically non-toxic (above 15 g/kg). Therefore, the data from our study indicate that hydroalcoholic extract of *C. officinalis* bark and *A. indica* seed are both moderately toxic, *C. officinalis* aqueous bark extract and *Garcinia lucida* hydroalcoholic seed extract are slightly toxic while all the other plant materials extracts are practically non-toxic. Notwithstanding that the results of this study corroborate with some of those previously reported on the plants tested on vertebrate models, further investigations are required under identical experimental conditions (in terms of preparation of the extracts) to determine the correlation between the LD50s in animal models and in *G. mellonella* in order to allow a better extrapolation of the results to humans. Finally, given the large number of larvae used in this study (1100 larvae), this model may be of great utility if it is standardized as a study model for toxicity and may avoid animal sacrifices.

## 4. Conclusion

The data presented in this study indicate that some plants such as Cinchona officinalis (aquous and hydroalcoholic bark extract), and hydroalcoholic seed extract of Azadirachta indica and Garcinia lucida induced a high toxic effect on *G. mellonella* larvae; while other plant materials such as water extract of *Moringa oleifera, Cymbopogon citratus* and *Vernonia amygdalina* leaves exerted low toxicity. The high variability of the lethal dose values from one plant to another demonstrates the good sensitivity of *G. mellonella* and allows us to conclude that *G. mellonella* can be used as a reliable and robust system model for the assessment of the toxicity of medicinal plants. However, further studies are needed to establish the exact correlation between LD50, LD90, LD100 values in *G. melonella* and in vertebrate models in order to facilitate reliable and precise extrapolations in humans.

## Acknowledgments

This study has been supported by the RUDN University strategic Academic Leadership Program

## Notes

### Competing Interest Statement

The authors have declared no competing interest.

## References

1. Bharadwaj, K. C., Gupta, T., Singh, R. M. 2018. Alkaloid group of Cinchona officinalis: structural, synthetic, and medicinal aspects. In Synthesis of Medicinal Agents from Plants,205–227.

2. Júnior, W. S. F., Cruz, M. P., Dos-Santos, L. L., Medeiros, M.F.T., 2012. Use and importance of quina (Cinchona spp.) and ipeca (Carapichea ipecacuanha (Brot.) L. Andersson): Plants for medicinal use from the 16th century to the present. J. Herb. Med. 2(4), 103–112.

3. Sylvie, D. D., Anatole, P. C., Cabral, B. P., Veronique, P.B., 2014. Comparison of in vitro antioxidant properties of extracts from three plants used for medical purpose in Cameroon: Acalypha racemosa, Garcinia lucida and Hymenocardia lyrata. Asian Pac J Trop Biomed. 4, S625–S632.

4. Guedje, N. M., Tadjouteu, F., Onana, J. M., Nga, E. N., & Ndoye, O., 2017. Garcinia lucida Vesque (Clusiaceae): from traditional uses to pharmacopeic monograph for an emerging local plant-based drug development. J. Appl. Biosci. 109, 10594–10608.

5. Sonfack, C. S., Nguelefack-Mbuyo, E. P., Kojom, J. J., Lappa, E. L., Peyembouo, F. P., Fofié, C. K., … Dongmo, A. B., 2021. The Aqueous Extract from the Stem Bark of Garcinia lucida Vesque (Clusiaceae) Exhibits Cardioprotective and Nephroprotective Effects in Adenine-Induced Chronic Kidney Disease in Rats. Evid Based Complement Alternat Med T, 2021:5581041

6. Guedje, N. M., and Fankap, R., 2001. Utilisations traditionnelles de Garcinia lucida et Garcinia kola (Clusiaceae) au Cameroun. Systematics and Geography of plants, 747–758.

7. Kuete, V., Dongfack, M. D., Mbaveng, A. T., L allemand, M. C., Van-Dufat, H. T., Wansi, J. D., … Wandji, J., 2010. Antimicrobial activity of the methanolic extract and compounds from the stem bark of Drypetes tessmanniana. Chin. J. Integr. Med. 16(4), 337–343.

8. Etame, R. M. E., Mouokeu, R. S., Poundeu, F. S. M., Voukeng, I. K., Cidjeu, C. L. P., Tiabou, A. T., … Etoa, F. X., 2019. Effect of fractioning on antibacterial activity of n-butanol fraction from Enantia chlorantha stem bark methanol extract. BMC Complement Altern Med. 19(1), 1–7.

9. Ohemu, T. L., Agunu, A., Chollom, S. C., Okwori, V. A., Dalen, D. G., & Olotu, P. N.,2018. Preliminary phytochemical screening and antiviral potential of methanol stem bark extract of Enantia chlorantha Oliver (Annonaceae) and Boswellia dalzielii Hutch (Burseraceae) against Newcastle disease in Ovo. European Journal of Medicinal Plants. 25(4), 1–8

10. Abike, T. O., Osuntokun, O. T., Modupe, A. O., Adenike, A. F., Atinuke, A. R., 2020. Antimicrobial Efficacy, Secondary Metabolite Constituents, Ligand Docking of Enantia chlorantha on Selected Multidrug Resistance Bacteria and Fungi. Journal of Advances in Biology & Biotechnology. 23(6), 17–32.

11. Olivier, D. K., Van Vuuren, S. F., & Moteetee, A. N., 2015. Annickia affinis and A. chlorantha (Enantia chlorantha)–a review of two closely related medicinal plants from tropical Africa. J Ethnopharmacol. 176, 438–462.

12. Arévalo-Híjar, L., Aguilar-Luis, M. Á., Caballero-García, S., Gonzáles-Soto, N., & Valle-Mendoza, D., 2018. Antibacterial and cytotoxic effects of Moringa oleifera (Moringa) and Azadirachta indica (Neem) methanolic extracts against strains of Enterococcus faecalis. Int J Dent, 25,1071676.

13. Baildya, N., Khan, A. A., Ghosh, N. N., Dutta, T., & Chattopadhyay, A. P., 2021. Screening of potential drug from Azadirachta Indica (Neem) extracts for SARS-CoV-2: An insight from molecular docking and MD-simulation studies. J. Mol. Struct. 1227, 129390.

14. Tofel, K. H., Kosma, P., Stähler, M., Adler, C., & Nukenine, E. N., 2017. Insecticidal products from Azadirachta indica and Plectranthus glandulosus growing in Cameroon for the protection of stored cowpea and maize against their major insect pests. Ind Crops Prod. 110, 58–64.

15. Meisyara, D., Krishanti, N. P. R. A., Zulfitri, A., Lestari, A. S., Tarmadi, D., Himmi, S. K., … & Ismayati, M., 2019. Biological activity of local plant extracts from Toba Region as insecticide. In IOP Conference Series: Earth and Environmental Science.374(1), 012006

16. Nagar, N., & Devra, V., 2018. Green synthesis and characterization of copper nanoparticles using Azadirachta indica leaves. Mater. Chem. Phys. 213, 44–51.

17. Vijayakumar, S., Divya, M., Vaseeharan, B., Ranjan, S., Kalaiselvi, V., Dasgupta, N., … & Durán-Lara, E. F., 2020. Biogenic Preparation and Characterization of ZnO Nanoparticles from Natural Polysaccharide Azadirachta indica. L. (neem gum) and its Clinical Implications. J. Clust. Sci. 1–11.

18. Abbas, S., Nasreen, S., Haroon, A., & Ashraf, M. A., 2020. Synhesis of Silver and Copper Nanoparticles from Plants and Application as Adsorbents for Naphthalene decontamination. Saudi J. Biol. Sci. 27(4), 1016–1023.

19. Chinnasamy, G., Chandrasekharan, S., Koh, T. W., & Bhatnagar, S., 2021. Synthesis, Characterization, Antibacterial and Wound Healing Efficacy of Silver Nanoparticles From Azadirachta indica. Front. microbiol. 12, 204.

20. Aderinola, T. A., Alashi, A. M., Nwachukwu, I. D., Fagbemi, T. N., Enujiugha, V. N., & Aluko, R. E., 2020. In vitro digestibility, structural and functional properties of Moringa oleifera seed proteins. Food Hydrocoll. 101, 105574.

21. Anwar, F., Latif, S., Ashraf, M., & Gilani, A. H., 2007. Moringa oleifera: a food plant with multiple medicinal uses. Phytotherapy Research: An International Journal Devoted to Pharmacological and Toxicological Evaluation of Natural Product Derivatives. 21(1), 17–25.

22. Ray, S. J., Wolf, T. J., & Mowa, C. N.,2015. Moringa oleifera and inflammation: a mini-review of its effects and mechanisms. In “I International Symposium on Moringa”. 1158, 317–330.

23. Arulselvan, P., Tan, W. S., Gothai, S., Muniandy, K., Fakurazi, S., Esa, N. M., …& Kumar, S. S., 2016. Anti-inflammatory potential of ethyl acetate fraction of Moringa oleifera in downregulating the NF-κB signaling pathway in lipopolysaccharide-stimulated macrophages. Molecules. 21(11), 1452.

24. Saleem, A., Saleem, M., & Akhtar, M. F., 2020. Antioxidant, anti-inflammatory and antiarthritic potential of Moringa oleifera Lam: An ethnomedicinal plant of Moringaceae family. S Afr J Bot. 128, 246–256.

25. Tonukari, N. J., Avwioroko, O. J., Ezedom, T., & Anigboro, A. A., 2015. Effect of preservation on two different varieties of Vernonia amygdalina Del.(bitter) leaves. Food Sci. Nutr. 6(07), 623.

26. Dumas, N. G. E., Anderson, N. T. Y., Godswill, N. N., Thiruvengadam, M., Ana-Maria, G., Ramona, P., … & Emmanuel, Y., 2020. Secondary metabolite contents and antimicrobial activity of leaf extracts reveal genetic variability of Vernonia amygdalina and Vernonia calvoana morphotypes. Biotechnol Appl Biochem.2020

27. Hasibuan, P. A. Z., Harahap, U., Sitorus, P., & Satria, D.,2020. The anticancer activities of Vernonia amygdalina Delile. Leaves on 4T1 breast cancer cells through phosphoinositide 3-kinase (PI3K) pathway. Heliyon, 6(7), e04449.

28. Joseph, J., Lim, V., Rahman, H. S., Othman, H. H., & Samad, N. A. (2020). Anti-cancer effects of Vernonia amygdalina: A systematic review. Trop J Pharm Res. 19(8), 1775–1784.

29. Yedjou, C. G., Sims, J. N., Njiki, S., Tsabang, N., Ogungbe, I. V., & Tchounwou, P. B., 2018. Vernonia amygdalina delile exhibits a potential for the treatment of acute promyelocytic leukemia. Glob J Adv Eng Technol Sci. 5(8), 1–9

30. Egbuonu, A. C. C., & Amadi, R. P., 2021. Ethanolic Extract of Ground Vernonia Amygdalina Stem Exhibited Potent Antibacterial Activity and Improved Hematological Bio-Functional Parameters in Normal and Monosodium Glutamate-Intoxicated Rats. Journal of Applied Sciences and Environmental Management, 25(3), 311–317.

31. Wang, W. T., Liao, S. F., Wu, Z. L., Chang, C. W., & Wu, J. Y., 2020. Simultaneous study of antioxidant activity, DNA protection and anti-inflammatory effect of Vernonia amygdalina leaves extracts. Plos one, 15(7), e0235717.

32. Alara, O. R., & Abdurahman, N. H. (2021). Vernonia amygdalina leaf and antioxidant potential. Academic Press; In Toxicology. 347–353.

33. Do, D. N., Nguyen, D. P., Phung, V. D., Le, X. T., Le, T. M., Do, V. M., … & Luu, X. C., 2021. Fractionating of Lemongrass (Cymbopogon citratus) Essential Oil by Vacuum Fractional Distillation. Processes, 9(4), 593.

34. Muala, W. C. B., Desobgo, Z. S. C., & Jong, N. E., 2021. Optimization of extraction conditions of phenolic compounds from Cymbopogon citratus and evaluation of phenolics and aroma profiles of extract. Heliyon, 7(4), e06744.

35. Hanaa, A. M., Sallam, Y. I., El-Leithy, A. S., & Aly, S. E., 2012. Lemongrass (Cymbopogon citratus) essential oil as affected by drying methods. Ann. Agric. Sci, 57(2), 113–116.

36. Koné, C., 2020. Plantes médicinales dans la prise en charge des infections urinaires (Doctoral dissertation, USTTB).

37. Ignasiak, K., & Maxwell, A., 2017. Galleria mellonella (greater wax moth) larvae as a model for antibiotic susceptibility testing and acute toxicity trials. BMC Res. Notes. 10(1), 1–8.

38. Megaw, J., Thompson, T. P., Lafferty, R. A., & Gilmore, B. F., 2015. *Galleria mellonella* as a novel in vivo model for assessment of the toxicity of 1-alkyl-3-methylimidazolium chloride ionic liquids. Chem. 139, 197–201.

39. Allegra, E., Titball, R. W., Carter, J., & Champion, O. L., 2018. Galleria mellonella larvae allow the discrimination of toxic and non-toxic chemicals. Chem. 198, 469–472.

40. Wijesinghe, G. K., Maia, F. C., de Oliveira, T. R., de Feiria, S. N. B., Joia, F., Barbosa, J. P., … & Höfling, J. F.,2020 Effect of Cinnamomum verum leaf essential oil on virulence factors of Candida species and determination of the in-vivo toxicity with Galleria mellonella model. Mem Inst Oswaldo Cruz. 115: e200349.

41. Moya-Andérico, L., Vukomanovic, M., del Mar Cendra, M., Segura-Feliu, M., Gil, V., José, A., & Torrents, E., 2021. Utility of Galleria mellonella larvae for evaluating nanoparticle toxicology. Chem. 266, 129235.

42. Ogungbemi, A. O., & van Gestel, C. A., 2018. Extrapolation of imidacloprid toxicity between soils by exposing Folsomia candida in soil pore water. Ecotoxicology. 27(8), 1107–1115.

43. Land, M. H., Toth, M. L., MacNair, L., Vanapalli, S. A., Lefever, T. W., Peters, E. N., & Bonn-Miller, M. O., 2020. Effect of Cannabidiol on the Long-Term Toxicity and Lifespan in the Preclinical Model Caenorhabditis elegans. Cannabis Cannabinoid Res. 2020

44. Zhang, Q., Hao, L., & Hong, Y., 2021. Exploring the multilevel effects of triclosan from development, reproduction to behavior using Drosophila melanogaster. Sci. Total Environ. 762, 144170.

45. Piatek, M., Sheehan, G., & Kavanagh, K., 2020. Utilising Galleria mellonella larvae for studying in vivo activity of conventional and novel antimicrobial agents. Pathog. Dis. 78(8), ftaa059.

46. Mbarga, M.J.A., Podoprigora, I. V., Davares, A. K., Razan, M., Das, M. S., & Senyagin, A. N.,2021. Antibacterial activity of grapefruit peel extracts and green-synthesized silver nanoparticles. Veterinary World. 14(5), 1330.

47. Mouafo, H. T., Tchuenchieu, A. D. K., Nguedjo, M. W., Edoun, F. L. E., Tchuente, B. R. T., & Medoua, G. N., 2021. In vitro antimicrobial activity of Millettia laurentii De Wild and Lophira alata Banks ex CF Gaertn on selected foodborne pathogens associated to gastroenteritis. Heliyon. 7(4), e06830.

48. Situmorang, P. C., Ilyas, S., Hutahaean, S., & Rosidah, R., 2020. Components and acute toxicity of nanoherbal haramonting (Rhodomyrtus tomentosa). J HerbMed Pharmacol. 10(1), 139–148.

49. Gaponenko Y, Mialdun A, Shevtsova V., 2020. Diffusion of Quinine with Ethanol as a Co-Solvent in Supercritical CO2. Molecules. 25(22), 5372.

50. Maurice M.I., 2014. “Pharmacognostical Profile of Selected Medicinal Plants”, in Handbook of African Medicinal Plants au. Boca Raton: CRC Press, Routledge Handbooks Online. https://www.routledgehandbooks.com/doi/10.1201/b16292-4. Retrieved:01 juin 2021,

51. ECHA (European Chemicals Agency): https://echa.europa.eu/registration-dossier/-/registered-dossier/10428/7/3/1; 2021. Retrieved: 01/06/2021

52. Braga, T. M., Rocha, L., Chung, T. Y., Oliveira, R. F., Pinho, C., Oliveira, A. I., … & Cruz, A., 2021. Azadirachta indica A. Juss In Vivo Toxicity—An Updated Review. Molecules, 26(2), 252.

53. Islas, J. F., Acosta, E., Zuca, G., Delgado-Gallegos, J. L., Moreno-Treviño, M. G., Escalante, B., & Moreno-Cuevas, J. E., 2020. An overview of Neem (Azadirachta indica) and its potential impact on health. J. Funct. Foods. 74, 104171.

54. Kale, M., Dhanokar, S., Aher, A., Gawali, S., & Dhikale, R., 2020. Azadirachtin: Natures Gift to Mankind. Curr Trends Pharm Pharm Chem. 2(4), 40–44

55. CCOHS (Canadian Centre for Occupational Health and Safety Act). What is a LD50 and LC50?. https://www.ccohs.ca/oshanswers/chemicals/ld50.html. Retrieved: 02/06/2021

